# lncRNA-WAL Promotes Aggressiveness of Triple-Negative Breast Cancer via inducing β-Catenin nuclear translocation

**DOI:** 10.1101/2022.09.06.506751

**Authors:** Hongyan Huang, Haiyun Jin, Rong Lei, Zhanghai He, Shishi He, Jiewen Chen, Phei Er Saw, Zhu Qiu, Guosheng Ren, Yan Nie

## Abstract

Because of its insensitive to existing radiotherapy, chemotherapy and targeted treatments, Triple-negative breast cancer (TNBC) remains a great challenge to overcome. More and more evidence has indicated abnormal wnt/β-catenin pathway activation in TNBC but not luminal or her2+ breast cancer, and lncRNAs play a key role in a variety of cancers. Through lncRNA microarray profiling between Activated and inactivated Wnt/β-catenin pathway of TNBC tissues, lnc-WAL (Wnt/β-catenin associated lncRNA; WAL) was selected as the top up-regulated lncRNA in Wnt/β-catenin pathway activation compared with the inactivation group. RIP-seq was analyzed between β-catenin and IgG groups of, where lnc-WAL could interact with β-catenin. Clinically, increased lnc-WAL in the TNBC tumor tissue was associated with shorter survival. lnc-WAL promoted the EMT, the ability of breast cancer stem cells (BCSC), proliferation, migration and invasion of TNBC cells. Mechanistically, lnc-WAL inhibited β-catenin protein degradation via Axin-mediated phosphorylation at serine 45. Subsequently, β-catenin was accumulated in nuclear and activated the target genes. Importantly, Wnt/β-catenin pathway activation stimulated the transcription of lnc-WAL. These results pointed to a master regulatory role of lnc-WAL/Axin/β-catenin in the malignant progression of TNBC. Our findings provide important clinical translational evidence that lnc-WAL maybe as potential therapeutic target against TNBC.

## Introduction

In all types of breast cancer, TNBC is a refractory molecular subtype(KL et al., 2020, RL et al., 2020). It is insensitive to current adjuvant therapy and survival rates are considerably hindered for this disease(WD et al., 2010), where the prognosis of TNBC is the poorest among breast cancer subtypes(Q et al., 2019). An effective therapeutic strategy is still lacking for TNBC, hence, screening for effective therapeutic targets for TNBC are urgently needed(X et al., 2019, C et al., 2017).

Research has revealed that the Wnt/β-catenin pathway was activated in TNBC(AI et al., 2010, P et al., 2013). In our study, we also found that the quantity of β-catenin in the nucleus was increased to 67% in TNBC patients. It is well known about the activation of the Wnt/β-catenin pathway that β-catenin was not phosphorylated by destruction complexes, and subsequently β-catenin was translocated to the nucleus and target genes were transcripted(T et al., 2017, Cell, 2020, J et al., 2012, F et al., 2019, Wei et al., 2020). In many cancers, including colorectal cancer, activation of the Wnt/β-catenin pathway is mainly caused by inactivating mutations of APC and β-catenin(Morin et al., 1997, Peignon et al., 2011). However, somatic mutations (mutations of APC and β-catenin) that activated the Wnt/β-catenin pathway were rare in TNBC, hence there must be other mechanism that were closed to the activation of the Wnt/β-catenin pathway in TNBC(Raisch et al., 2019, Moumen et al., 2013).

Long noncoding RNAs (lncRNAs) are an endogenous class of lncRNAs longer than 200-nucleotides that are not translated into proteins and are involved in diverse pathological processes including cancer, inflammation, and other human diseases(F and Cell, 2018, P et al., 2018, S et al., 2019, Li et al., 2020). In previous studies, lncRNAs could modulate various biological and pathological processes by targeting the Wnt/β-catenin signaling pathway(H et al., 2019, BH et al., 2019, Lu et al., 2017). lnc-LALR1 inhibited AXIN1 expression by recruiting CTCF to the AXIN1 promoter region and activated the Wnt/β-catenin signaling pathway(Xu et al., 2013). lnc-00673-v4 activated Wnt/β-catenin signaling by augmenting the interaction between DDX3 and CK1 and consequently increased the aggressiveness of lung adenocarcinoma(Guan et al., 2019). In our previous study, lnc-NKILA could directly interact with functional domains of signaling protein NF-κB/IκB and prevented the activation of NF-κB to suppress breast cancer metastasis(Liu et al., 2015).

Herein, we report a lnc-WAL that was aberrantly expressed and pivotal for the function of the Wnt/β-catenin pathway in TNBC. Our study showed that lnc-WAL specifically bound β-catenin and AXIN, which resulted in dephosphorylation of β-catenin and nuclear translocation, sequentially enhancing the Wnt/β-catenin pathway activation. In turn, the activation of Wnt/β-catenin signaling promoted the production of lnc-WAL, demonstrating positive feedback between the lnc-WAL and Wnt/β-catenin pathway. Therefore, lnc-WAL promoted the aggressiveness of TNBC via activating the Wnt/β-catenin signaling pathway.

## RESULTS

### Identification of Wnt/β-catenin pathway activation-associated lncRNAs in TNBC

The Wnt/β-catenin pathway was abnormally activated and the reason remains unclear in TNBC(Clevers and Nusse, 2012). We detected the location and expression β-catenin by IHC in our cohort BC tissues samples, nuclear and cytoplasm β-catenin was most prominent in TNBC, membrane β-catenin was observed predominantly in luminal A, luminal B and HER2+ tumors (Fig. 1A and 1B). consistent with original reports. In order to detect the clinical significance of its activation in TNBC, the location of β-catenin was detected by IHC in 189 TNBC patients’ samples. The location of β-catenin was used to divide these patients into activation (β-catenin^Cytoplasm+Nuclear^) and inactivation (β-catenin^Membrane^) subgroups (Fig. S1A and S1B). Survival analysis revealed that patients with the activation subgroup had a poorer overall survival (OS) and disease-free survival (DFS) than inactivation subgroup (Fig. S1C and S1D). These results indicated that the Wnt/β-catenin pathway maybe a key factor in TNBC progression.

**Figure 1.**
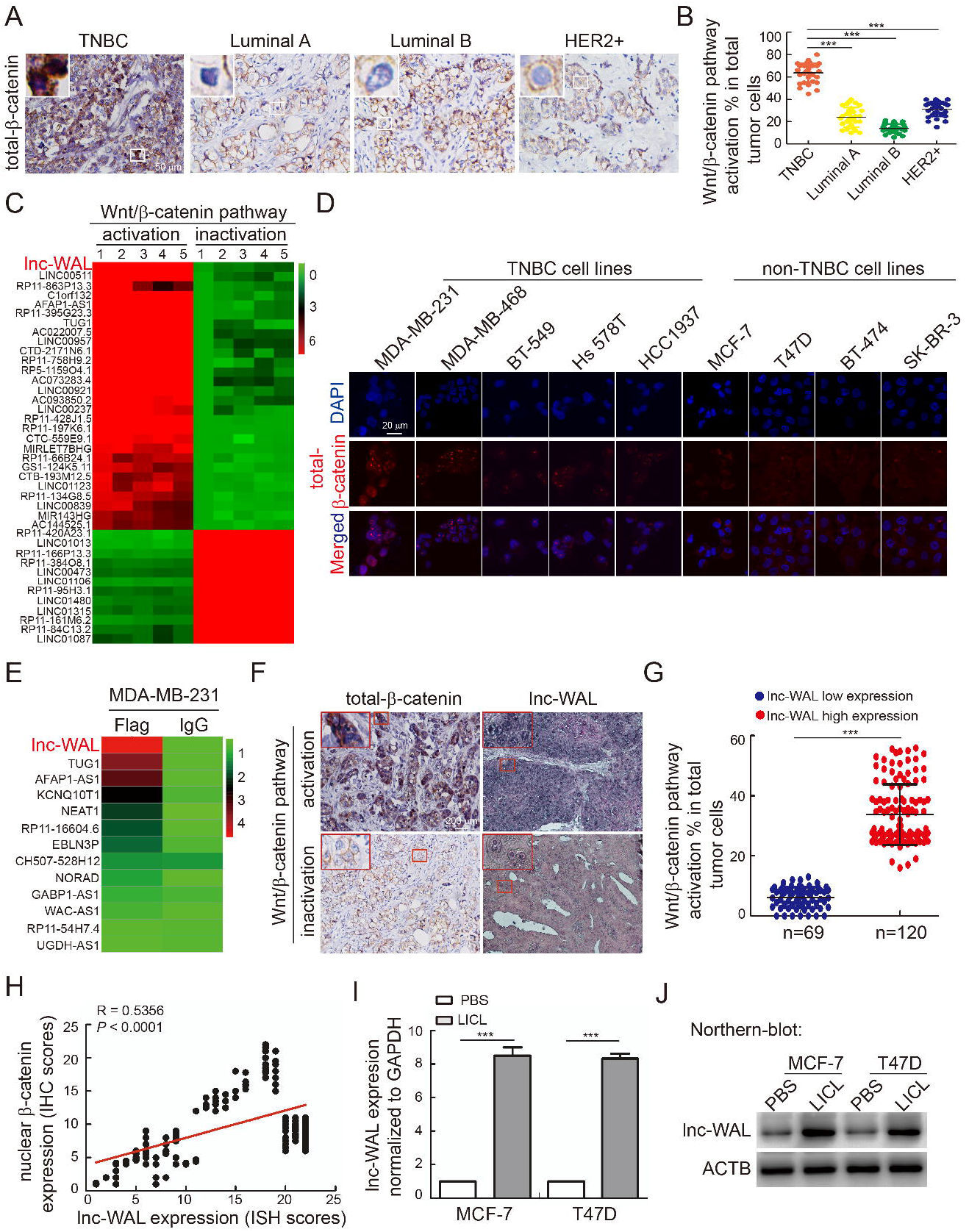
The identification of Wnt/β-catenin pathway associated lncRNAs. (A) The location of total-β-catenin was detected by IHC, it was located predominantly at the nuclear and cytoplasm in TNBC tissues, membrane total-β-catenin is most prominent in luminal A, luminal B and HER2+ BC tissues. The representative images of total-β-catenin location were shown in different molecular subtypes samples. Scale bar, 50 μm. (B) The percentage of nuclear and cytoplasm β-catenin in tumor cells of different molecular subtypes samples. (n = 30; mean ± SD; ***p < 0.001 by paired Student’s t tests) (C) LncRNAs expression profile in TNBC tissue samples with activation and inactivation of Wnt/β-catenin pathway by lncRNA microarray. (D) Immunofluorescent β-catenin staining showed the location of β-catenin in different subtypes breast cancer cells. (E) Heat map of lncRNAs that bound to β-catenin via Flag vs IgG group by RNA-seq in MDA-MB-231. (F) The expression of β-catenin by IHC and lnc-WAL by ISH in TNBC tissue samples with activation and inactivation of Wnt/β-catenin pathway. Scale bars, 200 μm. (G) Statistical chart of G showing that Quantification of Wnt/β-catenin pathway activation ratio in the lnc-WAL low expression group (n = 69) and the lnc-WAL high expression group (n = 120). (H) The correlation between the expression of lnc-WAL and the expression of nuclear β-catenin in 189 TNBC tissues. Pearson’s correlation coefficient r and p value were shown. (I and J) The expression of lnc-WAL can be stimulated by Wnt/β-catenin pathway agonist LICL by qRT-PCR and Northern blot.

In the TNBC, to further identify Wnt/β-catenin pathway activation-associated lncRNAs, we carried out one lncRNA tissues microarray to compare the difference among activation group (n=5) and inactivation group (n=5) in TNBC tissues. We screened out fifty-six lncRNAs were up-regulated and twenty-eight were down-regulated by more than fourfold (Table S1) and we identified lnc-WAL as the first up-regulated lncRNA in Wnt/β-catenin pathway activation group by the microarray profiling (Fig. 1C). To screen the lncRNAs that bound to β-catenin, we also detected the β-catenin location in cell lines by IF. Location of β-catenin prominently was in nuclear in the TNBC cells, compared to those others (Fig. 1D and Fig. S1E). Next, we screened the lncRNAs that bound to β-catenin between Flag/β-catenin and IgG in MDA-MB-231 by RIP-sequencing, where we also discovered that lnc-WAL was the strongest combination of β-catenin compared to IgG (Fig. 1E). we further verified the RIP-sequencing by RIP-qRT/PCR in MDA-MB-231(Fig. S1F). In clinical samples of TNBC patients, we examined the expression and location of β-catenin by IHC and the expression of lnc-WAL by ISH (Fig. 1F), we also performed statistical analysis about correlation between the β-catenin location and the lnc-WAL expression in 189 clinical TNBC samples, and found that the expression of lnc-WAL is higher in the activation samples than inactivation samples (Fig. 1G), and there was a positive relationship between the expression of lnc-WAL and β-catenin (Fig. 1H). In MCF-7 and T47D (Wnt/β-catenin pathway inactivation cell lines), qRT-PCR and NB arrays confirmed that lnc-WAL was up-regulated when the cells were disposed by LICL (Wnt/β-catenin pathway agonist) (Fig. 1I and 1J), and the expression of lnc-WAL in response to LICL was time- and dose-dependent (Fig. S1G-J). All these data above indicated that lnc-WAL was a Wnt/β-catenin pathway activation-associated lncRNA, and the functional characteristic of lnc-WAL has not been reported in previous research.

### lnc-WAL is highly expressed and predicts poorer prognosis of TNBC patients

Given that lnc-WAL was associated with Wnt/β-catenin signaling pathway activation in TNBC, we inspected the expression of lnc-WAL in different subtypes of breast cancer, where the level of lnc-WAL in TNBC was significantly elevated compared with Luminal and Her2+ breast cancer patients by qRT-PCR (Fig. 2A). Moreover, high levels of lnc-WAL were associated with Wnt/β-catenin signaling activation by qRT-PCR (Fig. 2B). Meanwhile, the lnc-WAL expression in TNBC cell lines was significantly elevated compared with the other subtypes breast cancer cell lines by qRT-PCR and NB (Fig. 2C and 2D).

**Figure 2.**
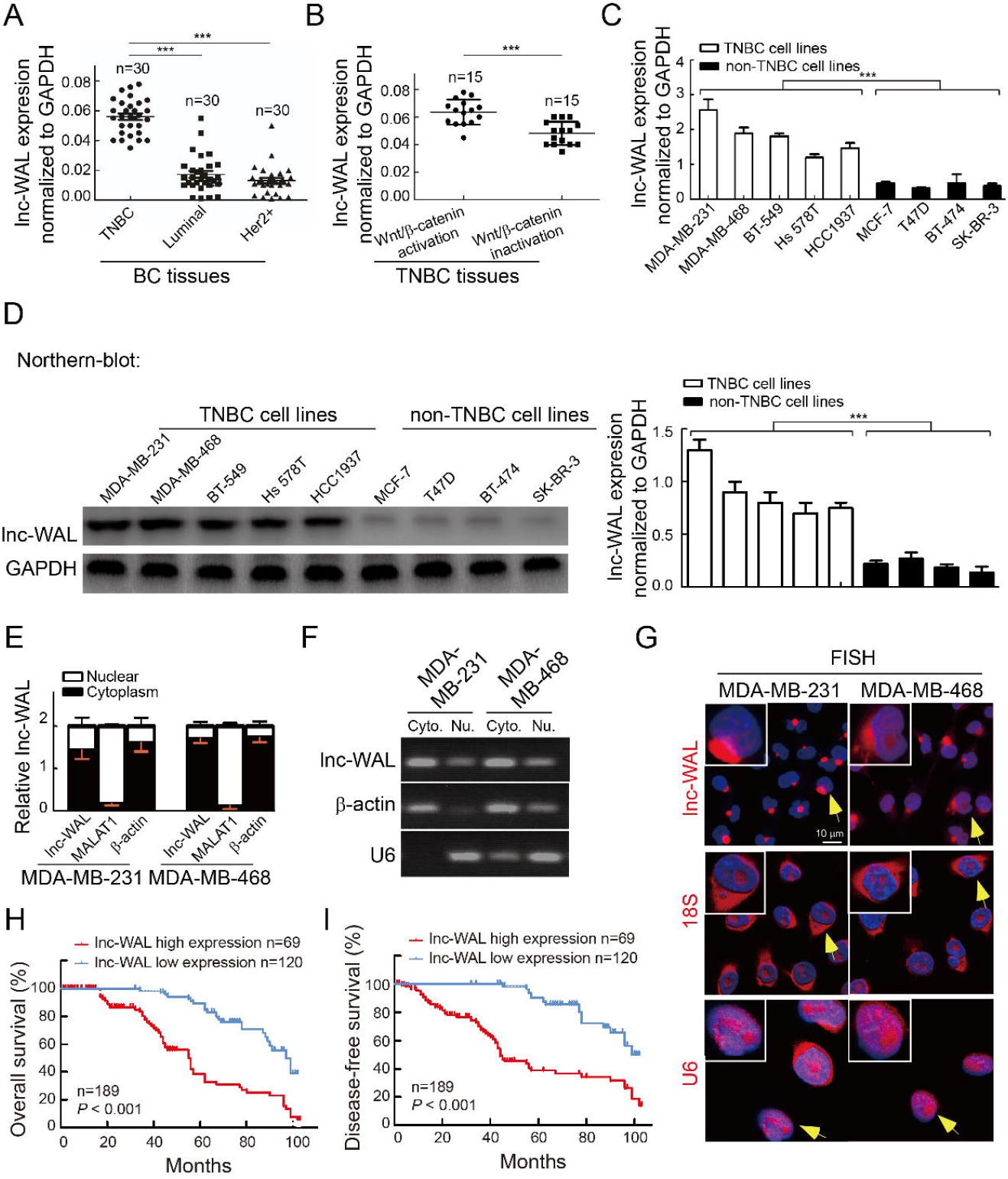
lnc-WAL is highly expressed and associated with poor prognosis in TNBC patients. (A) Expression of lnc-WAL was detected by qRT-PCR in different BC molecular subtypes tissue samples. (TNBC, n = 30; Luminal, n = 30; Her2+, n = 30). (B) Expression of lnc-WAL was detected by qRT-PCR in two groups: TNBC tissue samples of Wnt/β-catenin pathway activation (n = 15) and inactivation (n = 15). (C and D) The expression of lnc-WAL in different BC molecular subtypes cell lines by qRT-PCR and Northern blot. (E) The level of MALAT1 (nuclear control), β-actin (cytoplasmic control), as well as lnc-WAL were analyzed by qRT-PCR in nuclear and cytoplasmic (each bar represents the mean ± SD derived from three independent experiments). (F) RT-PCR showing lnc-WAL expression in the cytoplasm (Cyto.) and nucleus (Nu.) fractions of MDA-MB-231 and MDA-MB-468. MALAT1 was used as nuclear control, β-actin was used as cytoplasmic control. (G) Confocal FISH images showing cytoplasmic localization of lnc-WAL in TNBC cell lines MDA-MB-231 and MDA-MB-468, 18S and U6 as cytoplasm and nuclear control. Scale bars, 10 μm. (H) Kaplan-Meier survival curves analysis of TNBC patients with low and high lnc-WAL expression, the patients with high lnc-WAL had a poorer overall survival. (TNBC patients cohort, n = 189). (I) Kaplan-Meier survival curves analysis of TNBC patients with low and high lnc-WAL expression, the patients with high lnc-WAL had a poorer disease-free survival. (TNBC patients, n = 189).

In this study, after the nuclear and cytoplasmic fractionations of cells, qRT-PCR and RT-PCR showed that lnc-WAL mainly localized in the cytoplasm (Fig. 2E and 2F), which we also verified by RNA FISH, 18S and U6 as cytoplasm and nuclear control (Fig. 2G). The amplification of the lnc-WAL in the UCSC Genome Browser, where we performed 5′ and 3′ RACE assay and found that the full length of lnc-WAL was 4300 nt (Fig. S2A). These results suggest that the expression of lnc-WAL is higher in TNBC compared with other subtypes, especially in the TNBC with Wnt/β-catenin signaling activation.

We detected lnc-WAL using ISH in 189 TNBC tissue samples, and these patients were divided into lnc-WAL^High^ and lnc-WAL^Low^ subgroups by the staining intensity and positive percentage of lnc-WAL. We analyzed the relationship between lnc-WAL and the clinicopathological features of 189 TNBC tissue samples, and found that higher lnc-WAL expression was significantly relevant with tumor size, metastasis and local recurrence, Ki-67 level and Wnt/β-catenin pathway activation status, but was not associated with age, menopausal status, and histological grade (Table S2). Kaplan-Meier analysis indicated that patients with high lnc-WAL had a poorer OS and DFS than those with low lnc-WAL expression (*P* < 0.001, Fig. 2H and 2I). Univariate and multivariate Cox proportional hazard analyses showed that lnc-WAL (*P* < 0.001) was a poor independent prognostic factor for OS and DFS (Tables S3, 4). These results suggested that lnc-WAL levels are closely related to malignant development of TNBC, and can be used to predict the outcome of OS and DFS with TNBC.

Next, multiple datasets were retrieved. In the GEPIA datasets, higher lnc-WAL levels were associated with poorer OS (*P* = 0.03) and DFS (*P* = 0.0067) in BRCA (Fig. S2B and S2C). Strikingly, in multiple GEPIA cancer datasets, high lnc-WAL levels were statistically significantly associated with poor survival of liver, stomach and lung cancers (Fig. S2D - G). In the dataset of KM-plot TNBC, higher lnc-WAL levels had poorer DFS (*P* = 0.034). Additionally, in the dataset of stomach cancer KM-plot, higher lnc-WAL levels were associated with poorer OS (*P* = 7.6e-09, HR = 1.91, 95% CI = 1.53 to 2.38) (Fig. S2H and S2I). The above public available datasets were consistent with our survival data of BRCA samples.

### lnc-WAL promotes and maintains the malignant phenotype of TNBC cells

This part shows that lnc-WAL plays a key role in maintaining TNBC cells phenotype. The special characteristics of TNBC are EMT and the relatively high percentage of the CD44^high^/CD24^low^ cell population. To directly assess the role of lnc-WAL in EMT, siRNAs against lnc-WAL were transiently transfected into cells, where we found that siRNA against lnc-WAL induced the MET of MDA-MB-231 cells and largely reserved their mesenchymal morphology (Fig. 3A). WB revealed that lnc-WAL inhibited epithelial markers E-cad and promoted mesenchymal markers N-cad and Vimentin (Fig. S3A).

**Figure 3.**
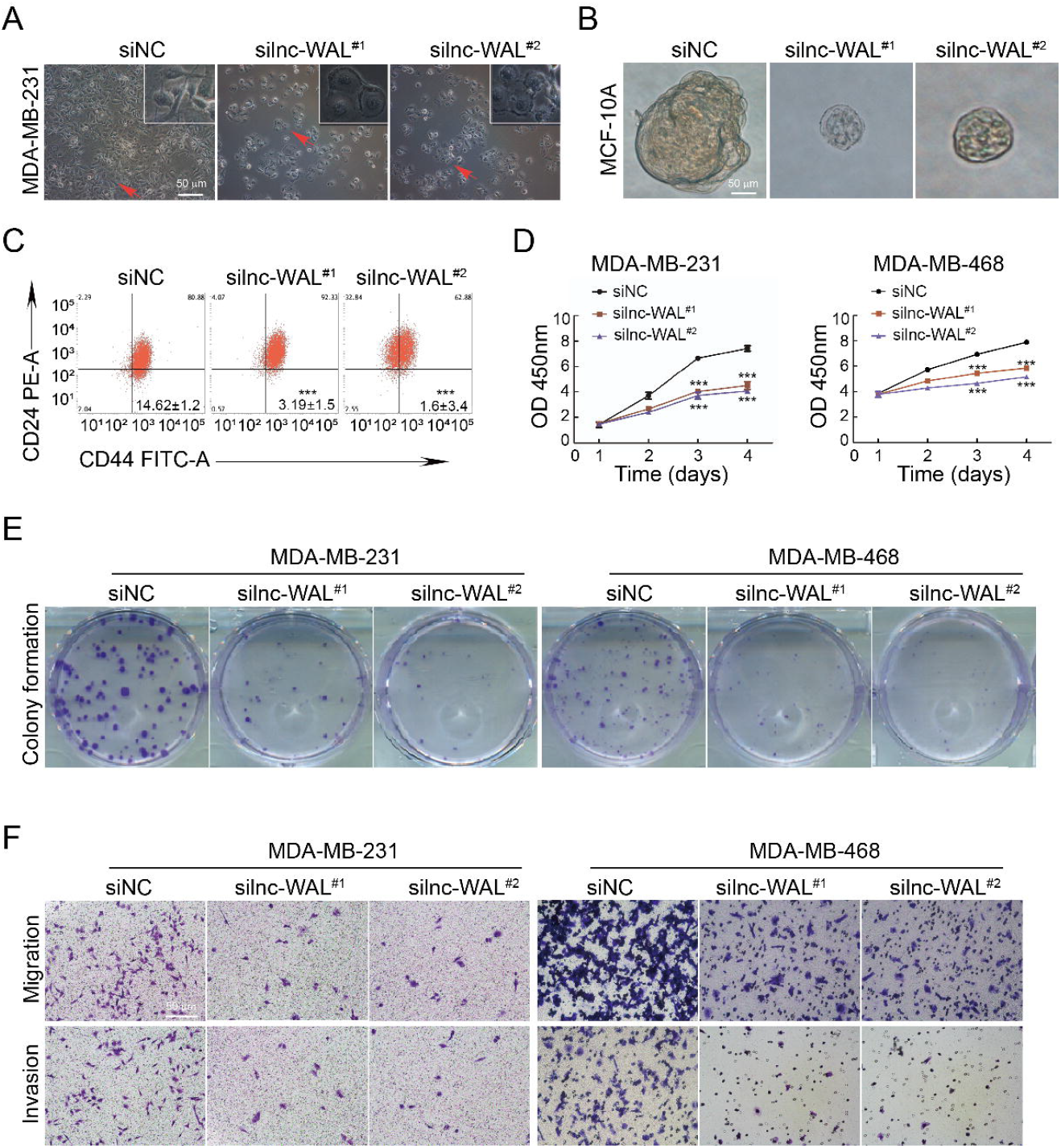
lnc-WAL induces EMT of TNBC cells and supports niche for BCSCs. (A) Representative images of MDA-MB-231 cell morphology after silencing lnc-WAL upon transfection with siRNAs. Scale bars, 50 μm. (B) Representative images of sphere formation in MCF-10A cells. Scale bars, 50 μm. (C) The percentage of CD44^+^CD24^-^ cells in the MCF-10A cells with or without silencing of lnc-WAL upon transfection with siRNAs by flow cytometry. (***p < 0.001 by paired Student’s t tests, mean ± SD). (D) Effects of lnc-WAL knockdown on proliferation in MDA-MB-231 and MDA-MB-468 cells by MTS assay. (E) Effects of lnc-WAL knockdown on colony formation in MDA-MB-231 and MDA-MB-468 cells. Scale bars, 50 μm. (F) Representative images of indicated migrating or invading cells analyzed by Matrigel-coated or noncoated Transwell assays in MDA-MB-231 and MDA-MB-468 cells, respectively. Scale bars, 50 μm.

To directly evaluate the role of lnc-WAL in breast cancer stem cells (BCSCs), we used immortalized human mammary epithelial cells MCF10A to detect the influence of lnc-WAL in the proportion of BCSCs(Clément et al., 2017), we found that lnc-WAL-depleted cells generated smaller and less mammospheres than siNC treated cells, while lnc-WAL could promote the mammosphere forming capacity of MCF10A cells (Fig. 3B and S3B). As expected, lnc-WAL could further increase BCSCs-enriched CD44^high^/CD24^low^ cell population by flow cytometry (Fig. 3C), and lnc-WAL increased CD44 and reduced CD24 expression detected by Western blot (Fig. S3C). Thus, lnc-WAL was a key gene in regulating BCSCs.

Here, we also found that lnc-WAL promoted cell proliferation, colony formation, migration and invasion. The above-mentioned functional experiments were carried out by using siRNAs. The proliferation indices (optical density values) of cells were significantly decreased in both silnc-WAL#^1^ and silnc-WAL#^2^ compared to siNC (Fig. 3D), lnc-WAL knockdown also inhibited colony formation capacity of cells at days 14 after siRNA transfection (Fig. 3E and S3D). Both cell migration and invasion were significantly reduced in both silnc-WAL^#1^ and silnc-WAL^#2^-treated compared to siNC-treated by migration and invasion assays (Fig. 3F and S3E). the knockdown efficiency of siRNAs targeting lnc-WAL was shown by qRT-PCR (Fig. S3F). These results indicate that lnc-WAL can promote the malignant progression of TNBC.

### lnc-WAL binds to β-catenin and inhibits β-catenin phosphorylation

lnc-WAL was screened out from those lncRNAs that bound to β-catenin by RIP-sequencing in MDA-MB-231. To verify whether lnc-WAL combined with β-catenin, we performed RIP-PCR and RNA pull-down. Our results indicated that lnc-WAL specifically interacted with β-catenin (Fig. 4A and 4B). In addition, lnc-WAL FISH and IF staining demonstrated that there was prominent colocalization between lnc-WAL and β-catenin in the cytoplasm (Fig. 4C).

**Figure 4.**
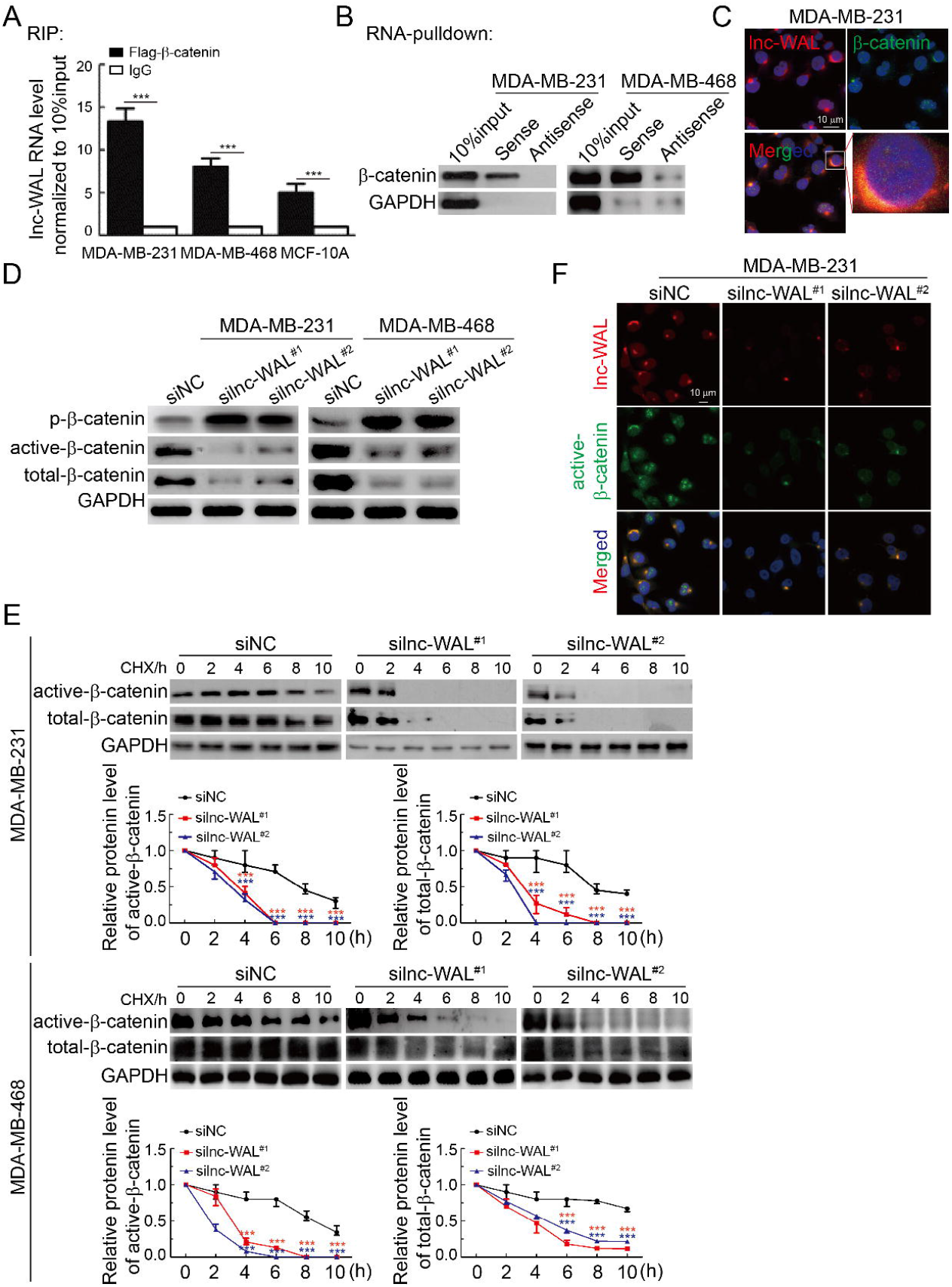
lnc-WAL binds to β-catenin and inhibits β-catenin phosphorylation. (A) RIP of β-catenin in MDA-MB-231, MDA-MB-468 and MCF-10A cells. lnc-WAL was retrieved by Flag-β-catenin IP, compared with IgG, as detected by qRT-PCR. ***p < 0.001. (B) RNA pulldown of lnc-WAL with total-β-catenin in MDA-MB-231 and MDA-MB-468 cells. RNA pulldown assay was performed using lnc-WAL or Antisense, followed by Western blot detection of total-β-catenin. (C) Confocal images showing the cytoplasmic colocalization of lnc-WAL and β-catenin in MDA-MB-231. Scale bars, 10 μm. (D) Silencing lnc-WAL upon transfection with siRNAs decreased the expression of active-β-catenin and total-β-catenin but promoted β-catenin phosphorylation by WB. (E) Protein degration assay with cycloheximide (CHX) 50 μ g/mL showing that the half-life of active-β-catenin and total-β-catenin were shortened after silencing lnc-WAL upon transfection with siRNAs. (F) A representative photo of active-β-catenin nuclear translocation, assayed by immunofluorescence and FISH confocal microscopy, in MDA-MB-231 cells with lnc-WAL silenced. Scale bars, 10 μm.

Translocation of β-catenin into nucleus is mediated by β-catenin phosphorylation via a destruction complex which consists of GSK3, AXIN, APC and CK1α(R and Cell, 2017, CS et al., 2018). To explore the role of lnc-WAL in participating Wnt/β-catenin signaling activation, after knocking down lnc-WAL, we found that β-catenin phosphorylation was increased and active, total-β-catenin was decreased (Fig. 4D and S4A). To further explore the effects of lnc-WAL on β-catenin, we detected the effects of lnc-WAL on the protein stability of β-catenin by half-life assay with or without CHX (0.1 μg/mL), where the half-life of β-catenin protein was calculated with and without silnc-WAL. It was found that active, total-β-catenin protein levels decreased by ~50% within 7 h and 7.5 h in the siNC-treated cells. However, active, total-β-catenin protein levels decreased by ~50% within 3 h and 3.5 h in the silnc-WAL-treated cells. The half-life of active, total-β-catenin protein was significantly shortened to 3 h and 3.5 h (Fig. 4E). These data indicated that lnc-WAL increased β-catenin protein stability and prolonged half-life. We further verified the colocalization between lnc-WAL and β-catenin protein in the cytoplasm, with subsequent knockdown of lnc-WAL which decreased β-catenin nuclear translocation by using FISH and immunofluorescence (Fig. 4F and S4B). we detected the Knockdown efficiency of siRNA by qRT-PCR (Fig. S4C). These results indicate that lnc-WAL activates Wnt/β-catenin pathway via specifically combining and inhibiting phosphorylation and subsequent nuclear translocation of β-catenin.

### Molecular mechanism of lnc-WAL inhibiting phosphorylation of β-catenin

It is well known that β-catenin protein is phosphorylated by kinase destruction complexes including GSK3, AXIN, APC and CK1α(CS et al., 2018). To identify potential kinases that involved lnc-WAL induced the inhibition of β-catenin phosphorylation, we performed rescue assays *in vitro* using different kinase inhibitors. We found that SKL2001 (AXIN inhibitor) could rescue the inhibition of β-catenin induced by silnc-WAL. However, CK1α inhibitor (IC261) and GSK3 inhibitor (LICL and AR2000) could not rescue the inhibiton of β-catenin (Fig. 5A and S5A and S5B). To further confirm that lnc-WAL inhibited β-catenin phosphorylation by kinase AXIN, we silenced AXIN by siRNA after silnc-WAL. Subsequently, lnc-WAL induced inhibition of β-catenin phosphorylation was abrogated when AXIN was silenced by two siRNAs (Fig. 5B and S5C and S5D). On the other hand, due to phosphorylation site of β-catenin at serine 45 (S45) by AXIN(DE et al., 2002), point mutations at the AXIN binding site (S45A) of β-catenin rescued silnc-WAL induced phosphorylation of β-catenin (Fig. 5C and S5E and S5F). The siRNA sequences in vitro were seen in Table S5.

**Figure 5.**
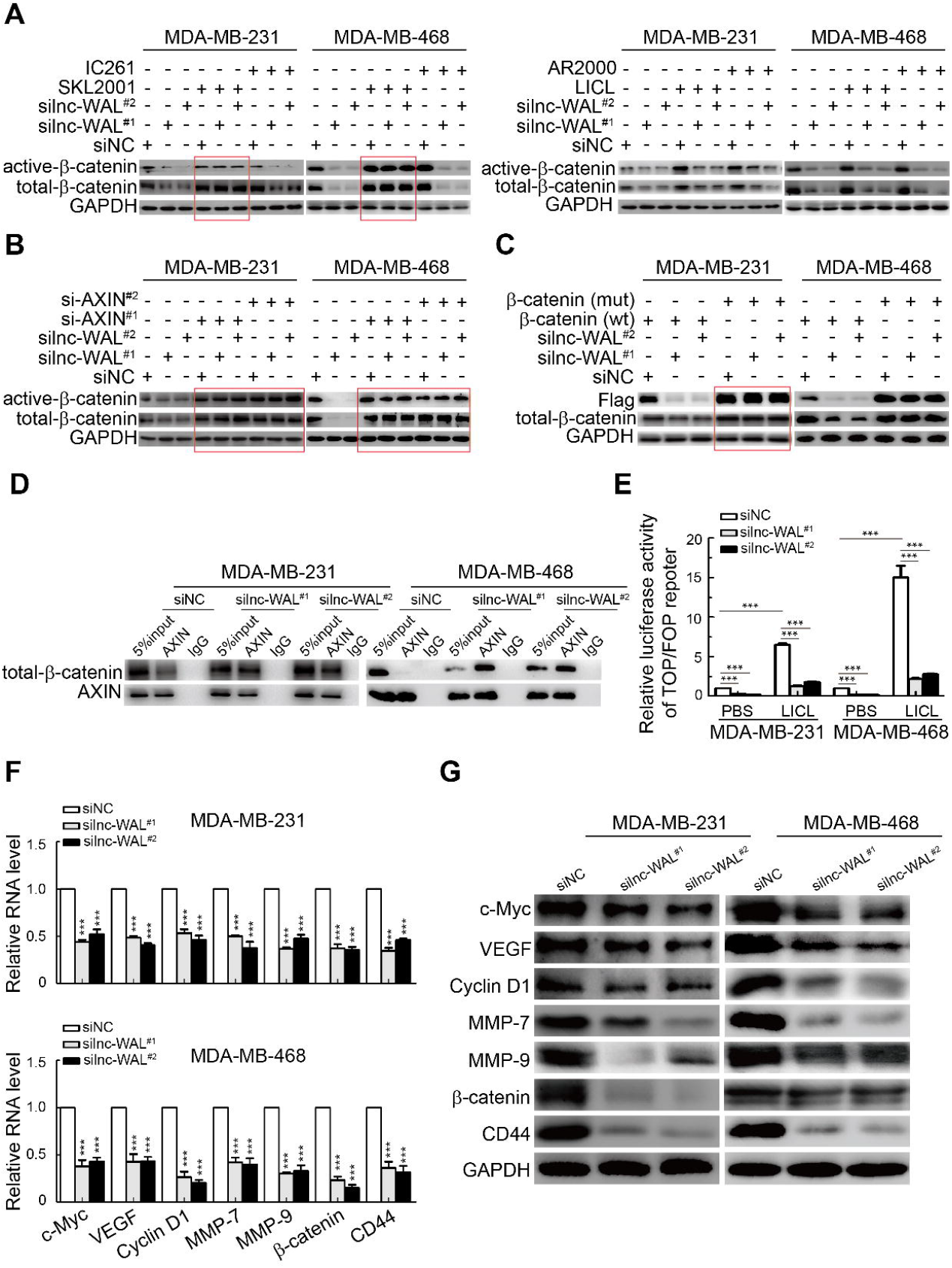
lnc-WAL inhibits β-catenin phosphorylation by physically obstructing AXIN. (A) Under the premise of silencing lnc-WAL with siRNAs, AXIN inhibitor SKL2001 rescued β-catenin (active-β-catenin and total-β-catenin) expression, but CK1α and GSK3 inhibitors could not induce this effect by WB in MDA-MB-231 and MDA-MB-468 cells. (B) Like AXIN inhibitor SKL2001, silencing AXIN by siRNAs rescued β-catenin expression (active-β-catenin and total-β-catenin) by WB in MDA-MB-231 and MDA-MB-468 cells. (C) After mutation of β-catenin S45 that β-catenin was phosphorylated by AXIN, the expression of active-β-catenin and total-β-catenin were rescued by WB in MDA-MB-231 and MDA-MB-468 cells. (D) Cell lysates of MDA-MB-231 and MDA-MB-468 cells with transient transfection of silnc-WAL or siNC were immunoprecipitated (IP) with anti-AXIN antibody followed by immunoblotting (IB). Knocking down lnc-WAL increased the interaction between AXIN and β-catenin. (E) β-catenin activity of LICL-treated MDA-MB-231 and MDA-MB-468 cells was examined by luciferase reporter assay (mean ± SD, ***p < 0.001). LICL was chosen due to optimal Wnt/β-catenin activator. (F and G) Knocking down lnc-WAL decreased the expression of Wnt/β-catenin pathway downstream genes by qRT-PCR and WB in MDA-MB-231 and MDA-MB-468 cells. GAPDH was the negative control.

To further understand whether lnc-WAL disturbs the binding of β-catenin and its kinase AXIN, we performed Co-IP and found that knockdown of lnc-WAL enhanced the interaction between β-catenin and AXIN (Fig. 5D). As it has been verified that binding between β-catenin and AXIN induces the phosphorylation of β-catenin and reduces nuclear accumulation of β-catenin(VS et al., 2012), we further explore whether the activation of Wnt/β-catenin pathway is through lnc-WAL to block the binding between β-catenin and AXIN. To this end, we knocked down lnc-WAL and found that it could activate Wnt/β-catenin pathway by luciferase report assay, LICL-treated cells were utilized as positive control (Fig. 5E). We examined whether lnc-WAL effects on downstream genes of the Wnt/β-catenin pathway, where the expression levels of c-myc, VEGF, cyclinD1, MMP-7, MMP-9, β-catenin and CD44 were decreased in cells where lnc-WAL was knocked down by qRT-PCR (Fig. 5F) and western-blot (Fig. 5G and S5G and S5H). Therefore, the results provide further evidence that lnc-WAL inhibits the phosphorylation of β-catenin via disturbing the binding of β-catenin and its kinase AXIN, which then induces active-β-catenin nuclear translocation.

### lnc-WAL promotes growth and distant metastasis of TNBC *in vivo*

To evaluate the therapeutic potential of targeting lnc-WAL *in vivo*, Indicated MDA-MB-231 cells (shvector, shlnc-WAL^#1^ and shlnc-WAL^#2^) were subcutaneously injected into 4 weeks old female nude mice, meanwhile, IWR-1 (the β-catenin inhibitor), which works by stabilizing AXIN was intraperitoneal injected (Fig. 6A). After 4 weeks injection, we found that both the volume and weight of tumors derived from lnc-WAL depleted group were clearly reduced compared to tumors derived from vector group. More importantly, the volume and weight of tumors derived from lnc-WAL depleted together with IWR-1 injection groups were further reduced compared to tumors derived from other groups (Fig. 6B - D). Moreover, hematoxylin eosin (HE) staining, ISH of lnc-WAL and IHC of β-catenin, Ki67 and immunofluorescence staining of TUNEL assay showed that lnc-WAL knockdown *in vivo* reduced cell proliferation, β-catenin activation and increased apoptosis of tumor cells respectively, while the effects were more significant in the groups (lnc-WAL depleted together with IWR-1) (Fig. 6E and S6A and S6B). We also examined the expression of lnc-WAL and target genes, we found that the levels of target genes were decreased in tumors samples derived from therapeutic groups compared to untreated group (Fig. 6F and S6C and S6D).

**Figure 6.**
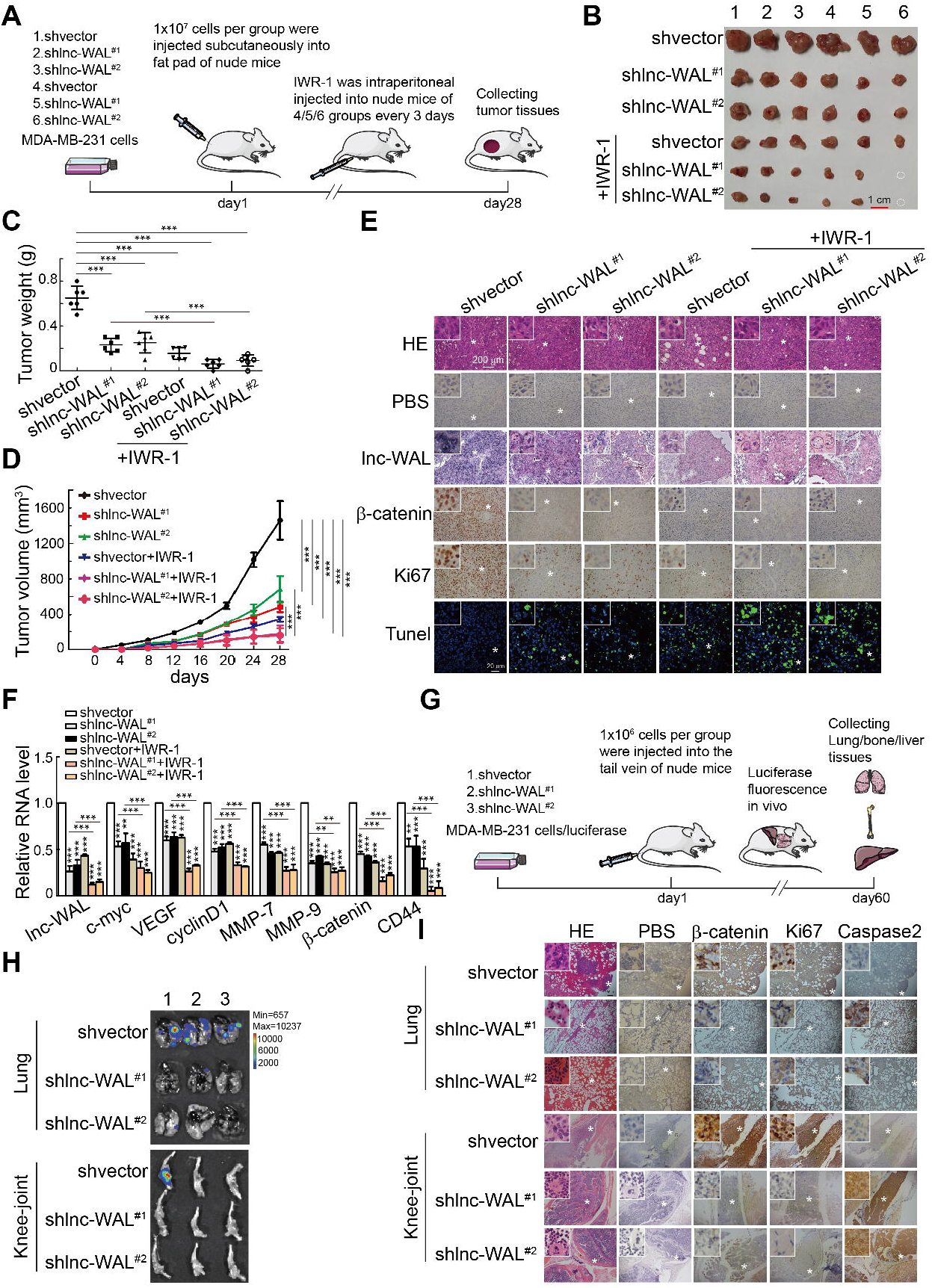
lnc-WAL promotes MDA-MB-231 cells growth and metastasis *in vivo*. (A) The process of constructing subcutaneous xenograft tumor in nude mice. (B) Indicated stable knockout MDA-MB-231 cells were injected into fat pads of xenograft nude mice with or without Wnt/β-catenin pathway inhibitor IWR-1 (0.8 μM). (C and D) Tumor weights and growth curves of different indicated MDA-MB-231 cells in nude mice. (mean ± SD, ***p < 0.001). (E) Representative in situ hybridization of lnc-WAL knockout efficiency for different groups in the paraffin-embedded sections of xenografts, β-catenin, ki-67 (the scale bar: 200 μm) and TUNEL assay staining (the scale bar: 20 μm) in the paraffin-embedded sections of xenografts. (F) Wnt/β-catenin pathway downstream genes c-myc, VEGF, cyclinD1, MMP-7, MMP-9, β-catenin and CD44 were detected by qRT-PCR in different groups. (G) The process of constructing metastasis xenograft tumor in nude mice. After injection of stably knocking out lnc-WAL MDA-MB-231 cells into the tail vein of nude mice at 60 days. (H) Bioluminescent images of lungs and knee-joints for each experimental group at 60 days. (I) HE staining of lung and knee-joint serial sections displayed metastatic nodules of the lungs and knee-joints. IHC staining of β-catenin, ki-67 and Caspase2 to investigate proliferation and apoptosis (the scale bar: 200 μm).

The metastasis rate of TNBC is up to 20%(RL et al., 2020). therefore, we established a mouse metastasis model using luciferase (Luc)-expressing MDA-MB-231 cells to further validate the promoting metastasis ability of lnc-WAL (Fig. 6G). We established stable knockout lnc-WAL by using two shRNA in Luc-expressing MDA-MB-231 cells, 10^6^ cells described above per mouse (shVector, shlnc-WAL^#*1*^ and shlnc-WAL^#*2*^) were intravenously injected into the 4 weeks old female nude mice (n = 3). After two months, all three mice developed obviously lung metastasis, and one of which also had obviously bone metastasis in knee joint. In contrast, the mice injected with shlnc-WAL^#*1*^ and shlnc-WAL^#*2*^ cells, did not develop any metastasis (Fig. 6H). The immunohistochemistry (IHC) staining of lung and knee joint serial sections also indicated that lnc-WAL promoted the tumor metastasis, as demonstrated by high lnc-WAL expression, more active-β-catenin expression, and more proliferation (Ki67), and less apoptosis (caspase 2) (Fig. 6I and S6E). All antibodies and reagent were seen in Table S6.

These results suggest that lnc-WAL is a promising therapeutic candidate that controls tumor growth and distant metastasis via regulating Wnt/β-catenin signaling pathway *in vivo*.

## Disscussion

Our study revealed the functional role of lnc-WAL in the malignant progress of TNBC via activating Wnt/β-catenin signaling, lnc-WAL was positively related with the progression of TNBC. At the molecular mechanism aspect, we found that lnc-WAL specifically perturbed the combination of AXIN with β-catenin at serine 45 to translocate signaling protein β-catenin into the nucleus. Under LICL stimulation, lnc-WAL expression was up-regulated in the MCF-7 and T47D. Therefore, under the condition of Wnt/β-catenin signaling hyperactivation, such as TNBC, lnc-WAL may function as a positive regulator of the Wnt/β-catenin signaling. The positive feedback loop was formed between Wnt/β-catenin signaling and lnc-WAL.

The Wnt/β-catenin signaling pathway plays a vital regulator in the development of all kinds of species. Whether in growth-related disease or different types of cancer, Wnt/β-catenin pathway is crucial, which suggests that it is a noticeable target for disease therapy(R and Cell, 2017, X et al., 2020, JH et al., 2019, W et al., 2019, KW et al., 2017). But, there are a lot of side effects if directly targeting Wnt/β-catenin pathway as cancer therapeutic strategy. Un-phosphorylation of β-catenin by destruction complexes (AXIN, GSK3 and CK1) and accumulation in the nucleus have been well recognized as a crucial part of Wnt/β-catenin signaling activation(P et al., 2019, EM et al., 2019). To explore molecular mechanism of Wnt/β-catenin signaling activation in TNBC, we performed a novel screening of the differentially expressed lncRNAs among activation and inactivation. Meanwhile, we searched for the lncRNA that binding to β-catenin by RIP-sequencing, and identified lnc-WAL as the most prominent lncRNA using the two methods mentioned above.

It has been reported that many lncRNAs regulate the Wnt/β-catenin signaling pathway. For example, lnc-CRNDE promoted colon cancer cells malignant evolvement by inhibiting miR-181a-5p expression, which indirectly influenced Wnt/β-catenin signaling(Han et al., 2017). By targeting miR-125b and miR-100, MIR100HG promoted cetuximab resistance via inhibiting negative regulators of Wnt/β-catenin pathway(Lu et al., 2017). In hepatobiliary carcinogenesis, activation of the Wnt/β-catenin pathway promoted the expression of lncRNA uc.158 and it may act as an endogenous competing lncRNA for the pro-apoptotic miR-193b, resulting in cancer cell survival(Fernández-Barrena et al., 2017). However, these articles just indicated the positive or negative relationship between lncRNA and Wnt/β-catenin pathway. Interestingly, lnc-00673-v4 acted as a scaffold to directly combine DDX3 with CK1, then promoted the aggressiveness of lung adenocarcinoma(Guan et al., 2019); lnc-CCAT1 could bind miR-204/211/148a/152 and inhibit their expression, leading to the upregulation of TCF4 and DNMT1, subsequently initiating the Wnt/β-catenin signaling in the breast cancer(T et al., 2019). Nevertheless, it is unclear whether there was some functional relationship among lncRNAs, signal protein β-catenin and kinase destruction complex (AXIN, GSK3 and CK1), especially in triple-negative breast cancer.

Unlike previously reported lncRNAs, lnc-WAL could directly bind signal protein β-catenin in the cytoplasm by RNA-pulldown, RIP and RNA-FISH experiments, where it inhibited β-catenin phosphorylation by AXIN kinase at serine 45 by utilizing kinase inhibition and point mutation assays. Co-IP assays demonstrated that lnc-WAL knockdown significantly enhanced the β-catenin and AXIN interactions. Finally, lnc-WAL activated Wnt/β-catenin pathway and promoted targeted gene production by luciferase report assays. Of note, we think it is not impossible that lnc-WAL activates Wnt/β-catenin signaling pathway through more than one mechanism, because β-catenin is phosphorylated by different kinases in different phosphorylation sites, in addition to the activation of β-catenin being a dynamic process from the cytoplasm to nucleus. Hence, it is beyond question that there will be different lncRNAs involved in different steps. Thus, whether there are any relationships among these lncRNAs that regulated Wnt/β-catenin pathway through the above putative mechanism also requires additional studies to be clarified.

Increasing proofs have demonstrated that differential expression of lncRNAs could be correlated with clinical prognosis in various cancers, which supports the possibility of lncRNA to be a targeted therapy in disease. Here, we found that lnc-WAL was significantly up-regulated in TNBC tissue samples, especially in Wnt/β-catenin signaling activation samples. The ectopic expression of lnc-WAL in TNBC tissue samples was closely associated with metastasis and relapse *in vitro* and *in vivo*. Our results indicated that TNBC patients of high lnc-WAL level have worse survival outcomes.

## Conclusion

In conclusion, lnc-WAL is a new reported lncRNA that regulates β-catenin phosphorylation by a kinase AXIN (Fig. S6F). This finding not only amplifies the regulatory mechanisms for Wnt/β-catenin signaling, but also draws attention to many lncRNAs which exerts their function by influencing the post-translational modification of signaling proteins. We conclude that lnc-WAL is indispensable in the Wnt/β-catenin signaling activation of TNBC and the regulatory role in this pathway is very crucial for the aggressive feature of TNBC. lnc-WAL could become a new therapeutic target for TNBC patients, there will less side effects compared with targeting Wnt/β-catenin signaling. We also will design more experiments and translational research to confirm the advantage of lnc-WAL in the treatment of TNBC.

## MATERIALS AND METHODS

### Patients and Tissue Samples

ISH staining for lnc-WAL and IHC staining for β-catenin were performed on the tissue samples of primary triple-negative breast cancer (189 cases) from Sun Yat-sen Memorial Hospital, Sun Yat-sen University (Guangzhou, China) and the First Affiliated Hospital of Chongqing Medical University (Chongqing, China) from May 2012 to March 2017. RNA levels of lnc-WAL were detected in the fresh frozen tissues of triple-negative (15 cases Wnt/β-catenin signaling activation and 15 cases Wnt/β-catenin signaling inactivation), Luminal (30 cases) and Her2^+^ (30 cases) samples from Sun Yat-sen Memorial Hospital (SYSMH), Sun Yat-sen University (Guangzhou, China). This study was approved by the Institutional Ethics Committee of the Sun Yat-sen Memorial Hospital, Sun Yat-sen University and First Affiliated Hospital of Chongqing Medical University.

### ISH, FISH, and IHC

The expression of lnc-WAL was detected in paraffin-embedded tissue sections and cell lines by using ISH and FISH arrays. ISH was performed using digoxigenin-labeled oligonucleotide lnc-WAL probe (TCAGCACTGTCATCATTACATT) (Takara, Japan) according to previous literature(Liu et al., 2015). For FISH, we used a FISH kit (Ribo^™^, Guangzhou, China), according to the manufacturer’s instructions. In the kit, 18S probe was used as cytoplasm control and U6 as nuclear control.

IHC was conducted as per the standard protocol according to literature(Su et al., 2018). The protein quantification was evaluated by two independent pathologists.

The ISH and IHC results were evaluated by two individuals in a blinded style using a quick scoring system from 0-12 by combing the staining intensity and the positive percentage, where the staining intensity was recorded on a scale of 0 (no staining); 1 (light staining); 2 (moderate staining); 3 (strong staining), the positive percentage was graded as follows: 0 (0%); 1 (<25%); 2 (25-50%); 3 (50-70%); 4 (>75%). The results were calculated as follows: staining intensity × positive percentage. This scoring system was based on previous literature(X et al., 2011).

### RIP and RNA sequencing

To explore those lncRNAs that bound to β-catenin, we established MDA-MB-231 cells with overexpressing β-catenin by transiently transfecting β-catenin/flag plasmid approximately 2 × 10^7^ cells were lysed by using IP lysis buffer (Invitrogen, 87787), and obtained lysates were incubated with Flag antibody and Dynabeads™ protein G (Invitrogen, 10004D) overnight at 4 °C with 360 degrees Celsius rotation and RNA was extracted using phenol-alcohol. RNA was detected by library construction and sequencing at Guangzhou GENE DENOVO Biotechnology Corporation.

### Cell nucleus/cytoplasm RNA fraction isolation

We used NE-PER^™^ Nuclear and Cytoplasmic Extraction Kit (Thermo, 78833) to detect the ratio of lnc-WAL in the nucleus and cytoplasm. Where the expression of lnc-WAL in Nuclear and Cytoplasm were detected by qRT-PCR. MALAT1 was used as nuclear positive control, and β-actin as the cytoplasm positive control. The associated gene primer sequences were listed in Table S7.

### RACE assay

RACE was conducted using SMARTer Kit (Takara, 634858) as the manufacturer’s instructions, where the detailed procedure was referenced to the literature(H et al., 2019). The associated 5’/3’ primer sequences were seen in Table S8.

### Sphere Formation Assay

The detailed process of the sphere formation experiment was performed as our previous instruction(Wei et al., 2015). MCF10A cells (800 cells/mL) were cultured in ultra-low adhesion plates (Corning) in serum-free DMEM-F12=1:1 (Gibco), including 4 mg/mL insulin (Sigma), 0.4% bovine serum albumin (Sigma), 20 ng/mL EGF (PeproTech), and B-27 (1:50, Gibco). After culturing for 7-10 days according to the design of the experiment, the diameter of mammospheres up to 75 μm were harvested, and three times independent experiments were conducted in each analysis.

### Cell migration and invasion assays

To explore the motor and metastasis abilities of TNBC cells, for imitating the cell barrier, we used Transwell chambers. For invasion assay, a layer of Matrigel (BD Bioscience) was placed above the chamber. Indicated cells were pretreated with silnc-WAL (RiboBio, Guangzhou) for 48 h, and was subsequently trypsinized and washed in PBS. Two thousand cells were seeded into the upper chambers in DMEM with 5% FBS, and the lower chambers were added into DMEM with 10% FBS. About 8 h, the chambers were collected and quantified by photographing in 3 random fields.

### Luciferase Reporter assay

Dual-luciferase reporter assay was used to detect the effect of lnc-WAL to β-catenin activity. β-catenin-Luc plus 5 ng pRL-TK-Renilla were transfected into the cells with Lipofectamine 3000 (Invitrogen, L3000-015) in 96-well plates. After 8 h, siRNAs were transfected into the cells with Lipofectamine 2000 (Invitrogen, 11668027). After another 8 h, dual-luciferase reporter assay was performed via the Dual-luciferase^®^ Reporter Assay System (Promega, E1910) as previously described. Luciferase activity was normalized to Renilla luciferase activity for each sample.

### Co-immunoprecipitation (Co-IP)

Co-IP assay was used to detect the interaction among different proteins. Indicated cells were washed in cold PBS and lysed with IP-lysis buffer (Thermo, 87787) with protease inhibitor, where the detailed processes were previously described.(Su et al., 2018) The lysates were immunoprecipitated with primary antibodies for incubation overnight at 4°C, and Dynabeads^™^ protein G (Invitrogen, 10004D) was added into the lysates for another 2 h at room temperature with 360 degrees Celsius rotation. After washing 3 times with the lysis buffer, the beads were adsorbed by a magnet and resuspended in protein loading buffer. Finally, the complex was analyzed by WB.

### Animal experiments

The animal experiments were approved by the Animal Care and Use Committee of Sun Yat-sen University. We purchased 4 weeks-old female Balb/c nude mice from GemPharmatech Corporation (Nanjing, China). We established stably knocking-out lnc-WAL in MDA-MB-231 cells by transducing lentivirus, and vector as negative control. 5 × 10^6^ cells in 100 μl PBS were injected into the third right mammary fat pads (6 nude mice every group). Once tumors reached 5 mm in diameter according to the standard modified formula Volume (mm^3^) = (length × height^2^)/2, IWR-1 (0.8 μM, Wnt/β-catenin pathway inhibitor) was used to deliver therapy to the tumors once per 3 days *in vivo*. After 32 days of treatment, the tumors were excised, tumor tissues were individually preserved for IHC, ISH, RNA and protein extraction. For metastasis model, we transducing luciferase lentivirus into MDA-MB-231, 1 × 10^6^ cells were mixed in 200 μl of PBS and tail intravenously injected. The lung, liver and bone were excised to examine the metastases using IVIS Lumina Imaging System.

### Statistical analysis

All data are are expressed as the mean ± SD values, P-values were calculated by two-tailed Stuent’ s t-test or one-way analysis of variance (ANOVA) followed by GraphPad Prism 5. Significant differences (*P⩽0.05; **P⩽0.01; ***P⩽0.001) are indicated. Each independent in vitro experiment included three independent replicates. Error bars reflect independent experiments.

## Acknowledgements

We would like to thank for the RIP-seq services and bioinformatics analysis provided by Genedenovo Biotechnology Co., Ltd. (Guangzhou, China). We are particularly grateful to Professor Erwei Song and Hai Hu from SYSMH for the supervision of study design.

## Authors’ Contributions

Conceptualization: H.Y.H., G.S.R., Y.N.; Methodology: H.Y.H., H.Y.J., R.L., Z.H.H., S.S.H., J.W.C., Z.Q.; Data analysis and interpretation: H. Y. H., H.Y.J., Y. N.; Writing and editing: H. Y. H., PE. S., Y. N.; Supervision: G.S.R., Y.N.; Funding: H.Y.H., Y.N. All authors read and approved the final manuscript.

## Ethics statement

The the research protocol was approved by the Sun Yat-sen Memorial Hospital of Yat-sen University. All animal experiments were approved by the Animal Care and Use Committee of Sun Yat-sen University (ethical approval number: L102012021040C).

## Supplemental Figure legends

**Figure S1. The identification of Wnt/β-catenin pathway associated lncRNAs.**

(A) The selective of Wnt/β-catenin pathway activation (red arrow) and inactivation by the location of β-catenin in triple-negative breast cancer tissues. Scale bar, 200 μm.

(B) Quantitation of (1A) (n = 189, ***p < 0.001 by paired Student’s t tests).

(C and D) Survival analysis revealed that patients in the Wnt/β-catenin pathway activation subgroup (n = 85) had a poorer OS and DFS than Wnt/β-catenin pathway inactivation subgroup (n = 104).

(E) Quantitation of (fig1E), the statistics of nuclear of β-catenin in different BC cell lines. (***p < 0.001 by paired Student’s t tests).

(F) lncRNAs of RNA immunoprecipitation (RIP)-sequencing bound with β-catenin as shown by RIP followed by qRT-PCR in MDA-MB-231. (***p < 0.001 by paired Student’s t tests).

(G) The expression of lnc-WAL in response to LICL in a time-dependent manner by qRT-PCR in MCF-7 and T47D cells.

(H) The expression of lnc-WAL in response to LICL in a dose-dependent manner by qRT-PCR in MCF-7 and T47D cells.

(I) The expression of lnc-WAL in response to LICL in a time-dependent manner by NB in MCF-7 and T47D cells.

(J) The expression of lnc-WAL in response to LICL in a dose-dependent manner by NB in MCF-7 and T47D cells.

**Figure S2. Association of lnc-WAL expression with the clinical outcome of multiple cancer patients.**

(A) Full length of lnc-WAL (ENST00000593427) predicted in the UCSC Genome Browser and measured lnc-WAL acquired by 5′ and 3′ RACE.

(B and C) Association of lnc-WAL expression with the overall survival and disease-free survival in all breast cancer patients (n=1068) in the GEPIA database.

(D) Association of lnc-WAL expression with the disease-free survival in TNBC patients (n=161) in KM plotter database. *P* value was based on two-sided long-rank test for Kaplan-Meier curves.

(E) Association of lnc-WAL expression with the overall survival in lung cancer patients (n=479) in the GEPIA database.

(F and G) Association of lnc-WAL expression with the overall survival and disease-free survival in liver cancer patients (n=361) in the GEPIA database.

(H) Association of lnc-WAL expression with the overall survival in stomach cancer patients (n=631) in the KM plotter database.

(I) Association of lnc-WAL expression with the disease-free survival in stomach cancer patients (n=385) in the GEPIA database.

**Figure S3. lnc-WAL promotes malignant progression of TNBC cells.**

(A) The EMT protein markers of MDA-MB-231 cells after silencing of lnc-WAL upon transfection with siRNAs were determined by Western blotting for E-cad, N-cad and Vimentin.

(B) Statistical chart of sphere formation number in MCF-10A cells. Mean ± SD, (***p < 0.001 compared to unsilenced MCF-10A cells by Student’s *t*-test).

(C) The expression of protein CD44 and CD24 after silencing of lnc-WAL upon transfection with siRNAs were determined by Western blotting in the MCF-10A cells.

(D) Quantitation of colony formation (fig3E). (***p < 0.001 by paired Student’s t tests, mean ± SD).

(E) Quantitation of migration and invasion (fig3F). (***p < 0.001 by paired Student’s t tests, mean ± SD).

**Figure S4. lnc-WAL promotes malignant progression of TNBC cells.**

(A) Quantitation of the expression of p-β-catenin, active-β-catenin and total-β-catenin (fig4D). (***p < 0.001 by paired Student’s t tests, mean ± SD).

(B) Quantitation of the lnc-WAL and β-catenin by IF (fig4F).

(C) Knockdown efficiency of lnc-WAL in the MDA-MB-231 and MDA-MB-468 cell lines. Cells were transiently transfected with negative control (siNC) or targeting lnc-WAL (silnc-WAL#^1^ and silnc-WAL#^2^). ***p < 0.001.

**Figure S5. lnc-WAL inhibits β-catenin phosphorylation by physically obstructing AXIN.**

(A and B) In the MDA-MB-231 and MDA-MB-468 cell lines, under the premise of silencing lnc-WAL with siRNAs, cells were treated by the inhibitors of AXIN, CK1α and GSK3, the statistical chart of β-catenin

(active-β-catenin and total-β-catenin) expression.

(C and D) In the MDA-MB-231 and MDA-MB-468 cell lines, cells were treated by silencing lnc-WAL and AXIN with siRNAs, then the statistical chart of β-catenin (active-β-catenin and total-β-catenin) expression.

(E and F) In the MDA-MB-231 and MDA-MB-468 cell lines, under the premise of silencing lnc-WAL with siRNAs, cells were transfected by the plasmid with mutation of β-catenin S45 that β-catenin was phosphorylated by AXIN, the statistical chart of β-catenin (active-β-catenin and total-β-catenin) expression.

(G and H) In the MDA-MB-231 and MDA-MB-468 cell lines, cells were treated by silencing lnc-WAL with siRNAs, then the statistical chart of the Wnt/β-catenin pathway downstream genes expression. All data are expressed as mean ± SD of three independent experiments. Student’s *f*-test and one-way ANOVA were used for comparison between two groups and multiple groups, respectively, ***p < 0.001.

**Figure S6. lnc-WAL promotes MDA-MB-231 cells growth and metastasis *in vivo*.**

(A) MDA-MB-231 cell line stably knocking out lnc-WAL was constructed by lentivirus and the expression of lnc-WAL was detected by qRT-PCR.

(B) The statistical chart of lnc-WAL by ISH, β-catenin and Ki67 expression by IHC, apoptosis tumor cells by TUNEL assay in tumors sections of *in vivo*.

(C and D) The expression of Wnt/β-catenin pathway downstream genes in different groups was detected by WB.

(E) The statistical chart of β-catenin, Ki67 and Caspase2 expression by IHC in lung and knee-joint tissues *in vivo*.

(F) Graphical illustration of the interaction between lnc-WAL and Wnt/β-catenin pathway in TNBC cells.

